# A Multitrait Locus Regulates Sarbecovirus Pathogenesis

**DOI:** 10.1101/2022.06.01.494461

**Authors:** Alexandra Schäfer, Sarah R. Leist, Lisa E. Gralinski, David R. Martinez, Emma S. Winkler, Kenichi Okuda, Padraig E. Hawkins, Kendra L Gully, Rachel L. Graham, D. Trevor Scobey, Timothy A. Bell, Pablo Hock, Ginger D. Shaw, Jennifer F. Loome, Emily A. Madden, Elizabeth Anderson, Victoria K. Baxter, Sharon A. Taft-Benz, Mark R. Zweigart, Samantha R. May, Stephanie Dong, Matthew Clark, Darla R. Miller, Rachel M Lynch, Mark T. Heise, Roland Tisch, Richard C. Boucher, Fernando Pardo Manuel de Villena, Stephanie A. Montgomery, Michael S. Diamond, Martin T. Ferris, Ralph S. Baric

**Affiliations:** Department of Epidemiology, University of North Carolina at Chapel Hill, Chapel Hill, NC, USA; Department of Medicine, Washington University School of Medicine, St. Louis, MO, USA; Department of Pathology & Immunology, Washington University School of Medicine, St. Louis, MO, USA; Marsico Lung Institute, University of North Carolina at Chapel Hill, Chapel Hill, NC, USA; Department of Genetics, University of North Carolina at Chapel Hill, Chapel Hill, NC, USA; Department of Microbiology and Immunology, University of North Carolina at Chapel Hill, Chapel Hill, NC, USA; Department of Pathology and Laboratory Medicine, University of North Carolina at Chapel Hill, Chapel Hill, NC; Lineberger Comprehensive Cancer Center, University of North Carolina at Chapel Hill, Chapel Hill, NC, USA; Rapidly Emerging Antiviral Drug Discovery Initiative, University of North Carolina, Chapel Hill NC, USA; Department of Molecular Microbiology, Washington University School of Medicine, St. Louis, MO, USA

## Abstract

Infectious diseases have shaped the human population genetic structure, and genetic variation influences the susceptibility to many viral diseases. However, a variety of challenges have made the implementation of traditional human Genome-wide Association Studies (GWAS) approaches to study these infectious outcomes challenging. In contrast, mouse models of infectious diseases provide an experimental control and precision, which facilitates analyses and mechanistic studies of the role of genetic variation on infection. Here we use a genetic mapping cross between two distinct Collaborative Cross mouse strains with respect to SARS-CoV disease outcomes. We find several loci control differential disease outcome for a variety of traits in the context of SARS-CoV infection. Importantly, we identify a locus on mouse Chromosome 9 that shows conserved synteny with a human GWAS locus for SARS-CoV-2 severe disease. We follow-up and confirm a role for this locus, and identify two candidate genes, *CCR9* and *CXCR6* that both play a key role in regulating the severity of SARS-CoV, SARS-CoV-2 and a distantly related bat sarbecovirus disease outcomes. As such we provide a template for using experimental mouse crosses to identify and characterize multitrait loci that regulate pathogenic infectious outcomes across species.

## Introduction

Studies support the hypothesis that natural host genetic variation contributes substantially to microbial susceptibility and disease severity (1, 2). However, the specific genes and alleles that regulate differential disease outcomes to respiratory virus infections remain largely unknown. The viral family *Coronaviridae* is comprised of several human and animal pathogens and at least five zoonotic CoV have emerged or rapidly expanded their geographic range in mammals in the 21^st^ century (3, 4). To date, the most significant human pathogens include the group 2B sarbecoviruses (SARS-CoV and SARS-CoV-2), which likely emerged from bats to cause human epidemic or pandemic outbreaks of severe acute respiratory infections, leading to substantial morbidity and mortality (5, 6). Many other high-risk group 2B SARS-like viruses including bat Sarbecoviruses (BtCoV), group 2C Middle East Respiratory coronavirus (MERS-CoV), and MERS-like bat CoVs also appear poised to cause future human epidemics or pandemics (7–9). Emerging sarbecoviruses vary widely in their ability to cause human and animal disease (10). The 2003 SARS-CoV strain caused ∼8,000 infections and ∼800 deaths leading to a 10% mortality rate, whereas SARS-CoV-2 has infected >500 million humans, leading to over 6.2 million deaths in an ongoing pandemic (11, 12). SARS-CoV and SARS-CoV-2 demonstrate a broad host range with the ability to cause variable disease outcomes that range from asymptomatic infection to death in other mammals, raising questions whether common underlying host genes contribute to disease across species (13). While the 2003 SARS-CoV had a higher mortality rate compared to SARS-CoV-2, COVID-19 outcomes can vary from asymptomatic to life-threatening acute respiratory disease syndrome and death in humans, supporting a documented role for viral and inter-host genetic control of disease severity (14–17). Mapping the underlying natural host gene variation that regulates susceptibility and disease severity after diverse sarbecovirus infections and across multiple species is expected to reveal common genetic loci that regulate pathogenic outcomes, inform threat and risk potential, lead to refined small animal models of human disease, and reveal novel targets for therapeutic interventions.

Mouse-adapted SARS-CoV and SARS-CoV-2 cause disease in mice by inducing acute respiratory distress syndrome (ARDS) and recapitulating several aspects of COVID-19 pathogenesis including age-dependent severe disease (18, 19). Mouse genetic reference populations (GRPs) also have been employed as highly relevant models of human disease and used to identify host susceptibility loci and genes, as well as gene networks and higher-level interactions that regulate phenotypic variation and disease severity. Among mouse GRPs, the Collaborative Cross (CC) has over 44 million single nucleotide polymorphisms (SNPs) and 4 million insertions and deletions (InDels), which segregate between the eight founder strains (20). In addition, several thousand novel variants (SNPs and small indels, as well as large deletions) exist within individual CC strains (21–24). As such, the CC population models human genetic diversity with both common and rare variants poised to impact a variety of phenotypes, including disease severity after acute viral infections, like influenza virus, flaviviruses, filoviruses, and coronaviruses (20, 25–31).

In this study, we leveraged the CC model to characterize the genetic susceptibility landscape of sarbecovirus infections in mice. We demonstrated that multiple loci regulate the host disease responses to this subgroup of coronaviruses. Specifically, we identified two genes *Ccr9* and *Cxcr6* within a multitrait QTL located on Chromosome 9 (Chr9), which have natural polymorphisms driving altered expression levels that correlate with SARS-CoV and SARS-CoV-2 disease severity in mice. This QTL shows conserved synteny with a locus in humans on Chr3 (which includes Ccr9 and Cxcr6) that was identified in COVID-19 human Genome-wide Association Studies (GWAS) predicting severe outcome and hospitalization. In total, several GWAS in humans have identified a common locus on Chr3 (3p21.31),that identified six genes associated with COVID-19 severity, susceptibility, respiratory failure and risk of hospitalization, and included *CCR9*, and *CXCR9* (14, 15, 32–34). In addition, across these human studies there was a high amount of genetic variation as well as different lead SNPs in this region, suggesting that multiple genes may regulate disease associations and severity. Our concordant susceptibility profiles of the CC011 and CC074 parent mouse lines, as well as *Ccr9* and *Cxcr6* deficient mice infected with SARS-CoV MA15, SARS-CoV-2 MA10 or BtCoV HKU3-SRBD MA demonstrate involvement of this locus in severe COVID-19 disease susceptibility across species and across different sarbecoviruses, as well as highlighting the utility of pre-emergence disease models. Our results also demonstrate that the CC mouse panel is well suited to identify and validate relevant susceptibility regions for other human infectious and chronic diseases, while providing focus for a deeper understanding of emerging sarbecovirus disease patterns in animal and human populations.

## Results

### Variable Disease Phenotypes after CC-RI Strain Infection

To extend our previous understanding of how genetic variation contributes to differential SARS-CoV outcomes, we studied groups of female mice from 5 different CC strains (CC005/TauUnc, CC011/Unc, CC020/TauUnc, CC068/TauUnc, and CC074/Unc) infected with 2003 SARS-CoV using our SARS-CoV MA15 model, which recapitulates many of the disease outcomes reported in humans (35). As these strains did not segregate alleles at previously identified SARS QTL, differences in disease would be due to uncharacterized genetic differences (27, 28). Infected mice were followed for weight loss and possible mortality through 4 days post-infection (**Figure 1A**). We observed a wide variation of weight loss across these strains. Due to divergent disease susceptibilities to SARS-CoV MA15 and narrow variation within a given strain, we focused on the disease resistant CC011/Unc (hereafter CC011, <5% weight loss) and the highly susceptible CC074/Unc strain (hereafter CC074, >15% weight loss). To assess disease progression in these strains in greater detail, we challenged additional groups of CC011 and CC74 mice with SARS-CoV MA15 (n=6, respectively). CC074 mice showed severe disease by 4 days post infection (dpi) marked by ∼80% mortality of ∼80% in 2 separate experiments, and a >15% loss of body weight. In contrast, CC011 had a maximal weight loss of 5% body weight and 100% survival (**Figure 1B and 1C**). Despite differences in weight loss, mortality, and congestion score, both strains had high levels of virus detectable in lungs at 2 and 4 dpi (**Figure 1D and E** respectively; mean ∼5×10^6^ PFU/lobe, CC011 n=6; CC074 n=1, 5 mice died). While confirming the original observation that these two strains exhibit distinct disease susceptibilities, these results also suggested that differences in host immune responses, rather than viral burden, drive the different disease outcomes in these strains (**Figure 1A**).

**Figure 1.**
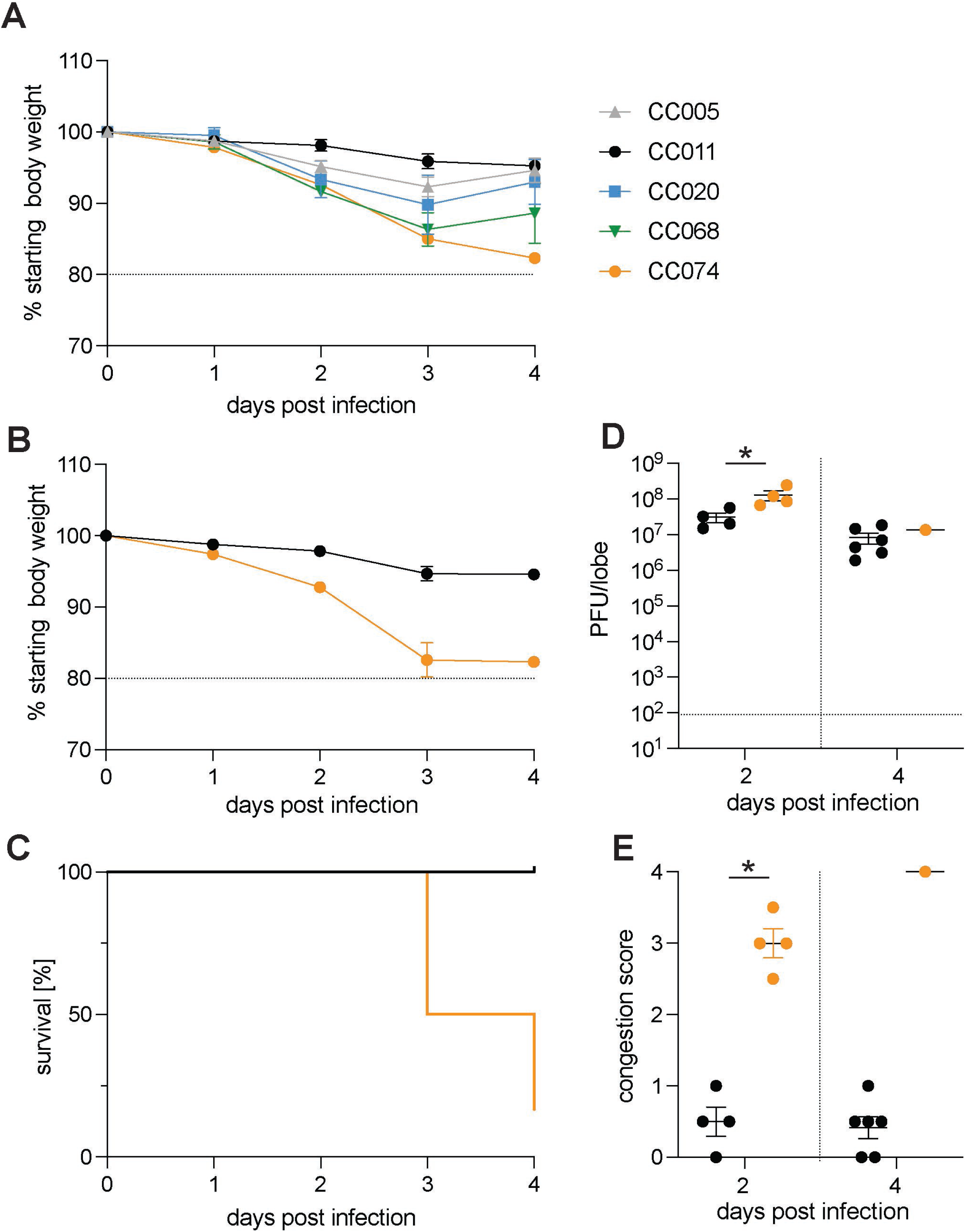
CC strains Demonstrate Different Susceptibility to SARS-CoV infection and disease. Age-matched female mice (n=4) of 5 different CC strains (CC005, CC011, CC020, CC068, and CC074) were infected with 1×10^4^ PFU and monitored for weight loss until 4 dpi. An additional group of age-matched CC011 and CC074 (both sexes) were infected with 1×10^4^ PFU and mice were monitored for disease progression until 4 dpi. A. Weight loss for CC strains CC005, CC011, CC020, CC068, and CC074 B. Weight loss of the two parental CC strains CC011 and CC074 C. Percentage survival of the two parental CC strains CC011 and CC074 D. Lung viral titer of the two parental CC strains CC011 and CC074 on 2 and 4 dpi E. Lung congestion score of the two parental CC strains CC011 and CC074 (female 10-12-week-old mice were infected with 1×10^4^ PFU SARS-CoV MA15; CC011: n = 4 for 2 dpi and n = 4 for 4 dpi; CC074: n =6 for 2 dpi and n=6 for 4 dpi respectively). Data were analyzed using Mann-Whitney test (lung titer and congestion scores), *p<0.05.

### SARS-CoV MA15 CC-F2 Screen

To identify the genetic basis of the observed disease outcomes and SARS-CoV pathogenesis, we generated a large F2 intercross between CC011 and CC074 (n=403 F2 mice, both males and females). F2 mice were inoculated intranasally at 9-12 weeks of age with 1×10^4^ PFU of SARS-CoV MA15. To expand our understanding of response to SARS-CoV infection, in addition to standard SARS-CoV-associated phenotypes including weight loss, viral burden, mortality, and lung congestion (**Figure 2A-D**), we also examined circulating immune cells as well as lung function (**Figure 2E-G**). Across these phenotypes, we observed an expanded range of disease responses in the F2 cross relative to their parent CC strains, (**Figure 2**, **Figure S1**). While a significant fraction of the F2 population (approximately 25%) died or were euthanized, this was diminished compared to CC074 (∼80% mortality).

**Figure 2.**
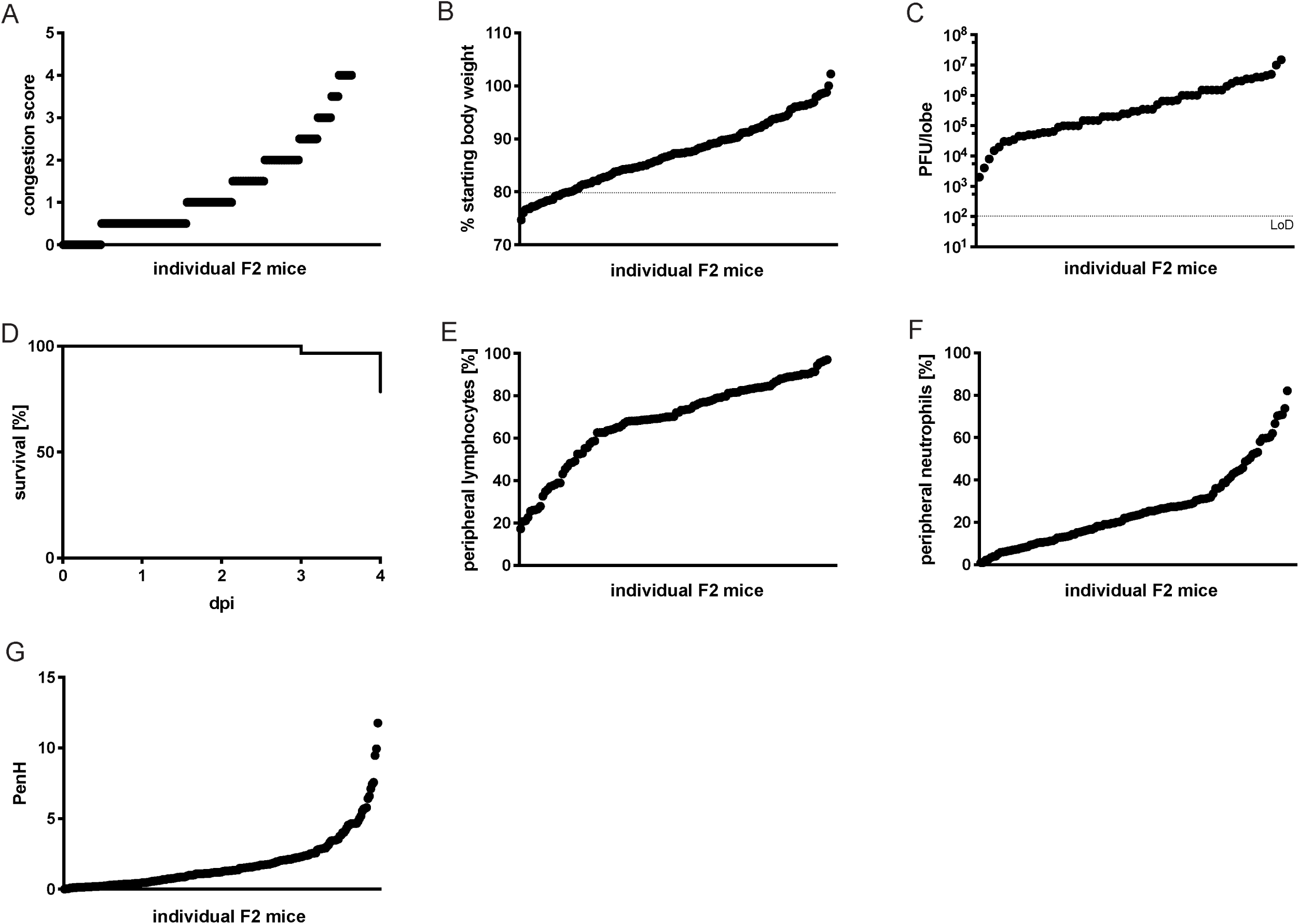
Disease phenotypes after SARS-CoV MA15 infection in the CC011xCC074-F2 Mice. 10-12-week-old CC011xCC074-F2 mice (n=403; 226 males, 177 females) were generated and infected with 1×10^4^ PFU of SARS-CoV MA15 and followed for 4 days to record clinical disease outcomes. A. Congestion score of CC011xCC074-F2 mice at 4dpi B. Weight loss of CC011xCC074-F2 mice C. Lung viral titer of CC011xCC074-F2 mice at 4dpi D. Percentage survival of CC011xCC074-F2 mice over the time of infection E. Percentage of peripheral blood lymphocytes of CC011xCC074-F2 mice at 4 dpi F. Percentage of peripheral blood neutrophils of CC011xCC074-F2 mice at 4 dpi G. PenH (airway resistance) of CC011xCC074-F2 mice at 2 dpi

Concurrent with viral challenge, the F2 animals were genotyped with the MiniMUGA array and QTL mapping in R/QTL conducted (36, 37). We identified a total of 2750 markers segregating between CC011 and CC074 among the autosomes and X-Chromosome. These markers were well distributed across the genome (23-48,228,264 nucleotides between markers, median = 509,769 nucleotides between markers). Using this genetic map, we identified a total of 9 genetic loci segregating in this F2 population for a variety of traits (Host response to SARS HrS24-28 at genome-wide p<0.05, HrS29-33 p<0.1), **Figure S1**, **Table 1**). Significant loci included a multitrait locus (Hrs26, Chr 9), as well as single loci associated with overall mortality (HrS24, Chr4), weight loss in males at 4 dpi (HsR25, Chr4), peripheral lymphocyte and neutrophil levels at 4 dpi (Hrs26, Chr11), and finally, altered pulmonary function (PenH) at 2 dpi (Hrs27, Chr15). The suggestive loci were identified in contributing to altered weight loss, mortality, and altered lung function phenotypes (**Table 1**).

**Table 1.**
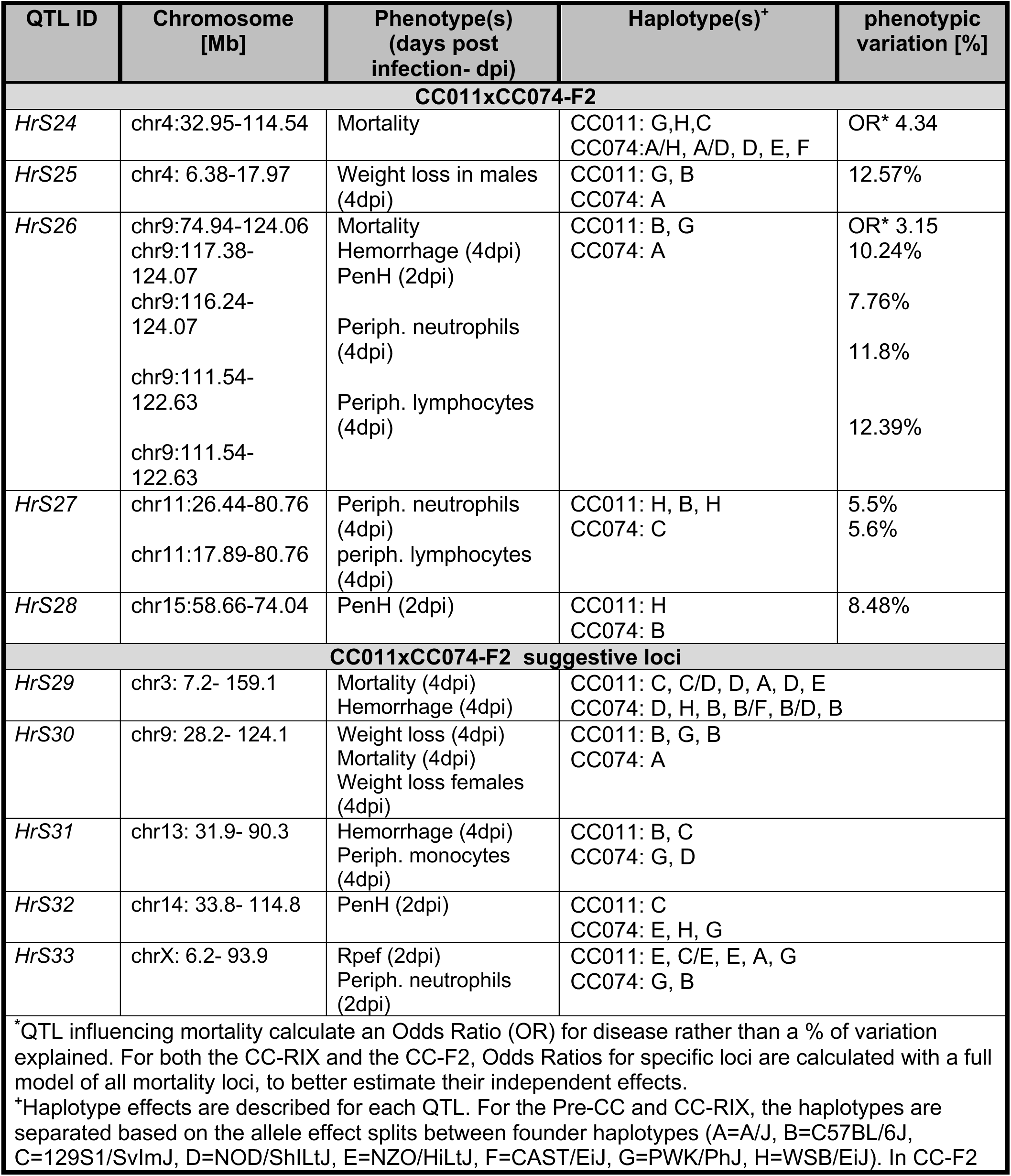
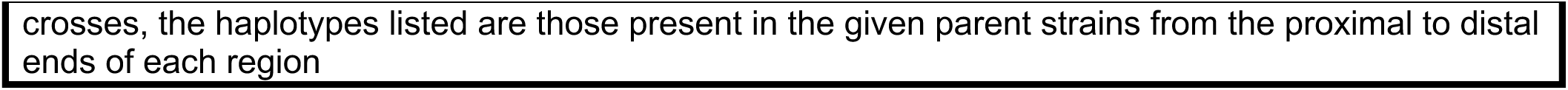
List of QTLs mapped with CC011xCC074-F2.

The multitrait HrS26 had a major impact on clinical disease (odds ratio (OR) of 3.15 for mortality), as well as lung pathology (10.24% of overall population variation) including lung function (7.76% of respiratory resistance (PenH) and immune cell inflammation (11.8% of neutrophils and 12.39% of lymphocyte variation) (**Figure S1, Table 1**). As such, it represents a locus of major effect driving disease differences between these strains following SARS-CoV MA15 challenge.

### CC011 and CC074 Disease Outcomes Following SARS-CoV-2 MA10 Infection

Concurrent with our ongoing analyses to phenotype SARS-CoV MA15 infection outcomes in CC parental and F2 strains, SARS-CoV-2 suddenly emerged. Although SARS-CoV-2 was genetically distinct with about 22% nucleotide differences across the genome as compared to SARS-CoV (6), the two viruses provided an opportunity to test whether similar host genes regulated pathogenesis across distinct sarbecoviruses. Accordingly, we infected CC011 and CC074 with the SARS-CoV-2 MA10 strain, which shares many of the COVID-19 disease phenotypes seen in human patients (**Figure 3**) (18). Importantly, the parental strains showed similar clinical disease phenotypes after SARS-CoV-2 MA10 infection (**Figure 3A-D**) and SARS-CoV MA15 (**Figure 1B-E**). Despite a small difference in weight loss and lung viral load between both virus strains, CC074 showed the same severe disease phenotypes with severe weight loss, lung discoloration scores, and 100% mortality by 4 dpi after infection with SARS-CoV-2 MA10 (**Figure 3C**). These results suggested that the QTL identified for SARS-CoV susceptibility, might also be relevant for SARS-CoV-2-induced disease.

**Figure 3.**
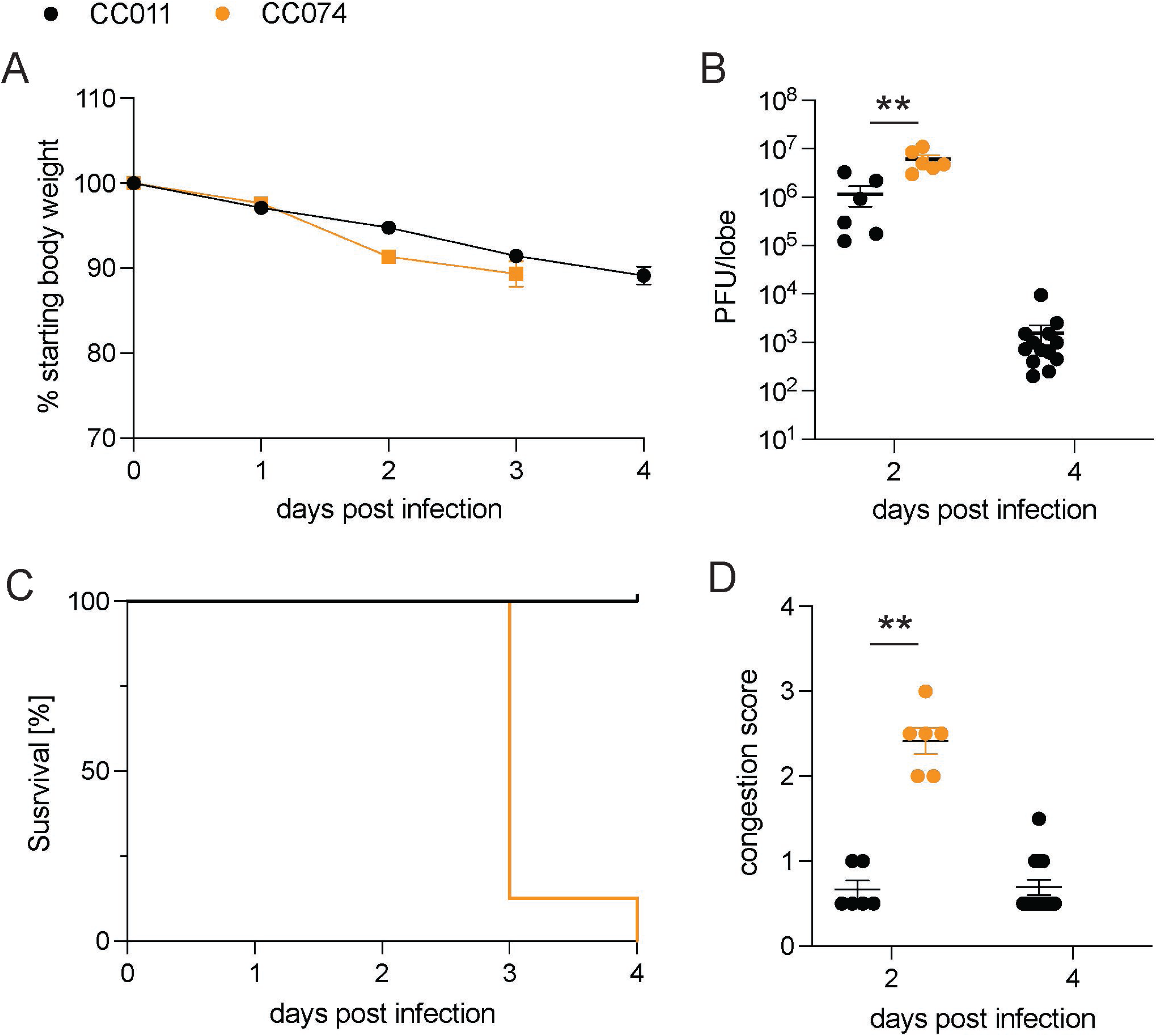
CC011 and CC074 Disease Outcomes after Mouse-adapted SARS-CoV-2 Infection. Groups of parental CC011 and CC074 mice were infected with 1×10^4^ PFU of SARS-CoV-2 MA10 and followed for several days for clinical disease outcomes. A. Weight loss of the two parental CC strains CC011 and CC074 B. Lung viral titer of the two parental CC strains CC011 and CC074 on 2 and 4 dpi C. Percentage survival of the two parental CC strains CC011 and CC074 D. Lung congestion score of the two parental CC strains CC011 and CC074 (female 10-12-week-old mice were infected with 1×10^4^ PFU SARS-CoV-2 MA10; CC011: n = 6 for 2 dpi and n = 13 for 4 dpi; CC074: n = 6 for 2 dpi and n =1 5 for 4 dpi respectively). Data were analyzed using 2-way ANOVA with Multiple comparison (weight and PenH), Log-rank (survival), and Mann-Whitney test (lung titer and congestion scores), *p<0.05, **p<0.005.

### Identification of a Multitrait QTL on Chr9

Recently, several reports have identified a common locus in humans on Chr3 (3p21.31) that is associated with COVID-19 severity, susceptibility, and risk of hospitalization (14, 15, 32–34). This locus, encompassing genes such as *SLC6A20*, *LZTFL1*, *FYCO1*, *XCR1, CXCR6*, and *CCR9,* shows conserved synteny with the proximal region of the multitrait locus mapped on Chr9 in mice (HsR26 and HsR30) (**Figure 4A, Table 1**). In addition, the studies in humans confirmed a high degree of polymorphism in this area on Chr3. We examined the sequence differences between CC011 and CC074 in this region of conserved synteny, where CC011 possessed a PWK/PhJ haplotype and CC074 had an A/J haplotype, and detected several SNPs and polymorphisms (**Table 2**) (38). *Ccr9* and *Fyco1* have missense SNPs in coding regions and all other SNPs were in regulatory regions such as the 5’- and 3’- UTR, at splice sites, and within introns, pointing towards differential expression of these target genes (**Table 2**). Similar findings and altered gene expression levels have been reported in the human GWAS, transcriptome-wide association study (TWAS), and epigenetic studies involving these gene sets (15, 16, 39–41).

**Figure 4.**
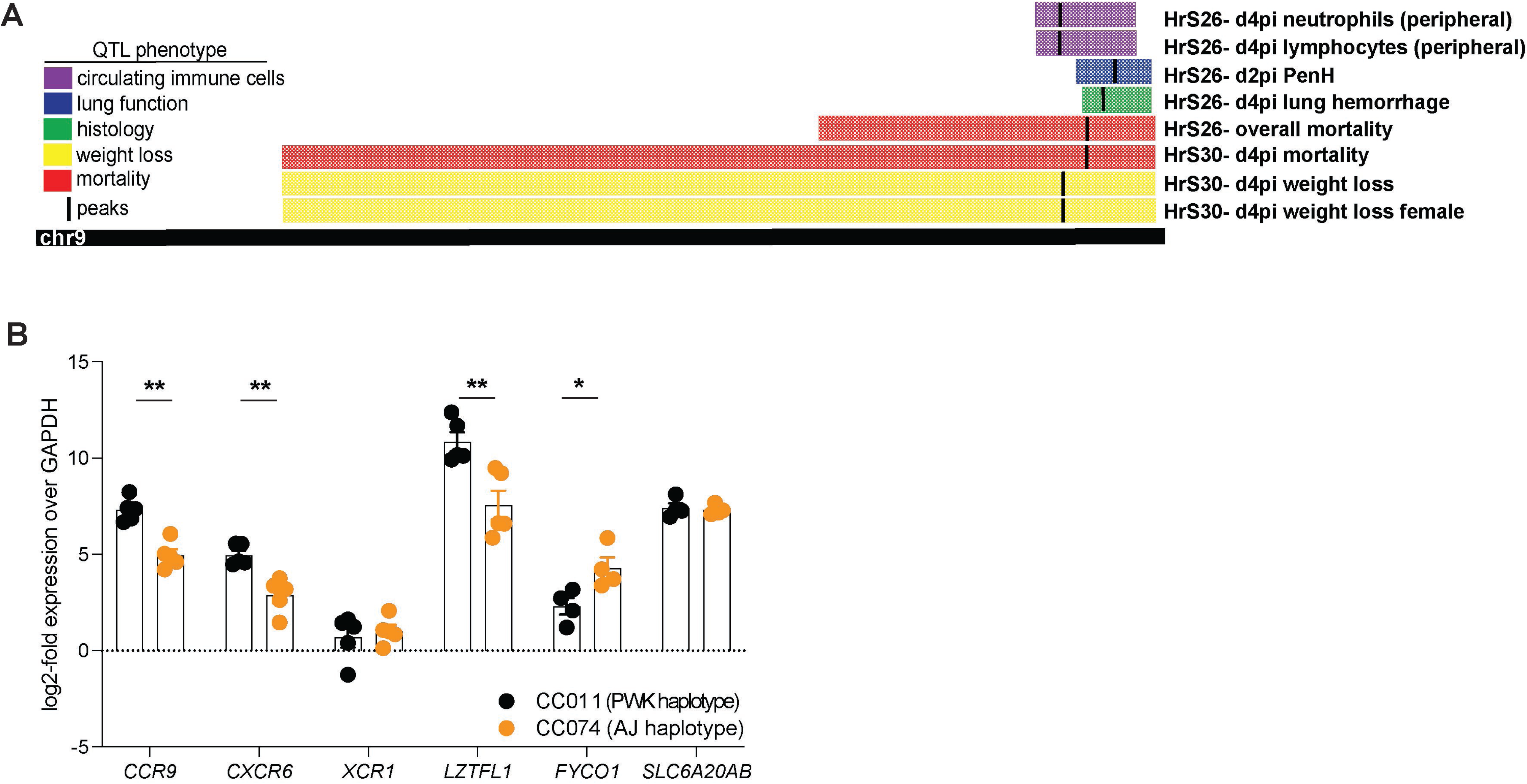
Identification of a Multitrait QTL on Chr9. Each of the individual F2 mice were genotyped using MiniMuga and QTL mapping conducted by testing the strength of association between each F2 mouse’s phenotype and their genotypes at each marker. A. A quantitative multitrait locus with major effect was identified on Chr9 (74.9–124 Mb), which affected mortality, weight loss, lung congestion, lung function, and peripheral hematology. CC011/Unc has a C57BL/6/PWK haplotype and CC074/Unc has an A/J / PWK haplotype in this QTL region. B. Expression levels of *Ccr9*, *Cxcr6, Xcr1, Lztfl1, Fyco1*, and *Slc6a20a/b* of SARS-CoV-2 MA10-infected CC011 and CC074 mice at 4 dpi, determined by quantitative RT-PCR, normalized to *Gapdh* (female 10-12-week-old mice were infected with 1×10^4^ PFU SARS-CoV-2 MA10; CC011: n = 5, CC074: n = 5) Data were analyzed using Mann-Whitney test, *p<0.05, **p<0.005.

**Table 2.** SNPs in CC011 and CC074 possible target genes in the Chr9 multitrait QTL (38) CC011xCC074-F2: Multi-Trait Locus on Chr9 Polymorphism

| variants | SCL6A20A/B | LZTFL1 | CCR9 | FYCO1 | CXCR6 | XCR1 |
| --- | --- | --- | --- | --- | --- | --- |
| missense | 8 |  | 3 | 20 |  |  |
| 5'UTR |  | 1 | 12 | 7 |  | 2 |
| 3'UTR | 7 | 33 | 57 | 729 | 3 | 34 |
| intron |  | 139 | 699 | 424 | 34 | 36 |
| splicing |  | 2 | 3 | 4 |  |  |

Consequently, gene expression profiles of *Ccr*9, *Cxcr6, Xcr1, Lztf1, Fyco1,* and *Slc6a20a/b* were measured in the lungs of infected CC011 and CC074. Significantly reduced expression of *Ccr9*, *Cxcr6*, and *Lztf1* was observed in highly susceptible CC074 mice (p<0.005), as compared to the resistant CC011 mice. In contrast, expression of *Fyco1* was significantly upregulated (p<0.05) in CC074 compared to the CC011 mice, while levels of *Xcr1* and *Slc6a20a/b* expression were not significantly different (**Figure 4B**). We took advantage of RNAseq data from unperturbed CC mice that contained either the protective (PWK) or susceptible (A/J) haplotype at this locus to determine the expression of these 6 genes (**Table S1**). We found significant differences in expression of *Lztf1* (increased in A/J), *Fyco1* (decreased in A/J), and *Slc6a20a/b* (increased in A/J), as well as a trend for *Xcr1* (increased in A/J). Together, these data suggest a complex cis-regulatory architecture of these genes at baseline along with induction in the context of infection.

### *Ccr9* Regulates Sarbecovirus Infection and Pathogenesis *in vivo*

The sarbecoviruses are divided into three phylogenetic clades designated clade Ia (SARS-CoV), clade II (HKU3) and clade Ib (SARS-CoV-2) (**Figure 5A**). To initially examine the impact of differences in gene expression have on virus disease phenotypes, we focused on *Ccr9* and *Cxcr6* which were differentially expressed by the respective mouse strains (**Figure 4B**).

**Figure 5.**
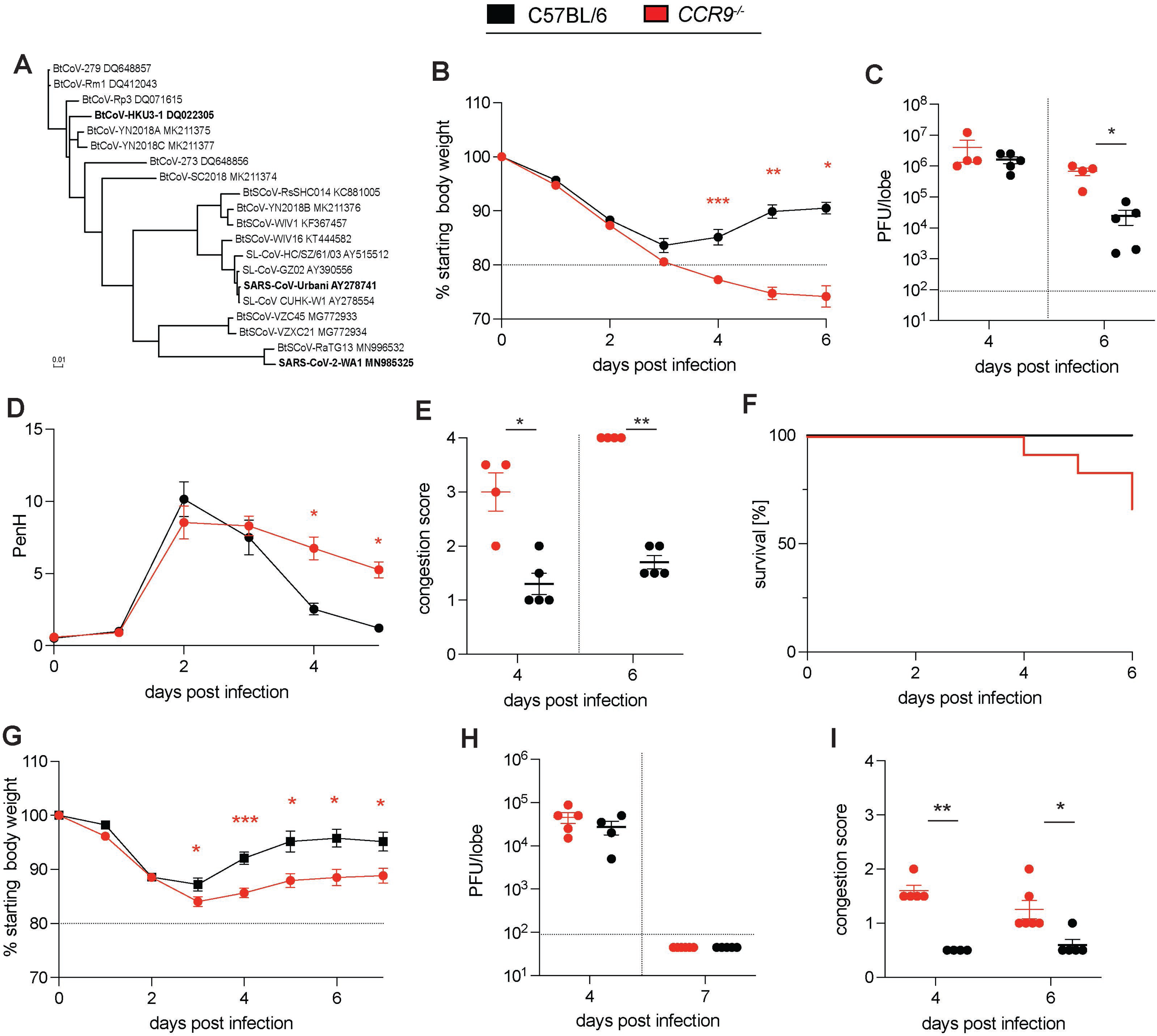
Validation of *Ccr9* as susceptibility gene during SARS-CoV MA15 and HKU3-SRBD MA infection. To validate *CCR9* as a susceptibility gene during SARS-CoV infection, groups of age-matched *CCR9^-/-^* mice were infected with 1×10^5^ PFU SARS-CoV MA15 and HKU3-SRBD MA and followed for several days for disease outcomes. The Spike protein sequences of selected Sarbecoviruses were aligned and phylogenetically compared. Sequences were aligned using free end gaps with the Blosum62 cost matrix, and the tree was constructed using the neighbor-joining method with a Jukes-Cantor genetic distance model based on the multiple sequence alignment in Geneious Prime. The GenBank accession numbers for each genome sequence are shown. The tree was then output and visualized using EvolView. A. Phylogenetic Tree of sarbecoviruses. The Spike protein sequences of selected Sarbecoviruses were aligned and phylogenetically compared. Sequences were aligned using free end gaps with the Blosum62 cost matrix, and the tree was constructed using the neighbor-joining method with a Jukes-Cantor genetic distance model based on the multiple sequence alignment in Geneious Prime. The GenBank accession numbers for each genome sequence are shown. The tree was then output and visualized using EvolView (Bold indicates viruses tested). B. Weight loss of *CCR9^-/-^* mice and C57BL/6NJ control mice C. Lung viral titer of *CCR9^-/-^* mice and C57BL/6NJ control mice on 4 and 6 dpi D. PenH of *CCR9^-/-^* mice and C57BL/6NJ control mice E. Congestion of *CCR9^-/-^* mice and C57BL/6NJ control mice on 4 and 6 dpi F. Percentage survival of *CCR9^-/-^* mice and C57BL/6NJ control mice G. Weight loss of *CCR9^-/-^* mice and C57BL/6NJ control mice H. Lung viral titer of *CCR9^-/-^* mice and C57BL/6NJ control mice on 4 and 7 dpi I. Lung congestion of *CCR9^-/-^* mice and C57BL/6NJ control mice on 4 and 7 dpi (15-18-week-old mice were infected with 1×10^5^ PFU SARS-CoV-2 MA10; C57BL/6NJ: n = 4 for 4dpi and n = 5 for d6pi; *CCR9^-/-^*: n = 5 for 4 dpi and n = 6 for 6 dpi respectively). Data were analyzed using Mann-Whitney test, *p<0.05, **p<0.005 (15-18-week-old mice were infected with 1×10^5^ PFU HKU3-SRBD MA; C57BL/6NJ: n = 5 for 4 dpi and n = 4 for 6 dpi; *CCR9^-/-^*: n = 5 for 4 dpi and n = 6 for 6 dpi respectively). Data were analyzed using 2-way ANOVA with Multiple comparison (weight and PenH), Log-rank (survival), and Mann-Whitney test (viral titer, congestion score), *p<0.05, **p<0.005, ***p<0.0005.

Knockout mice have the engineered construct introduced in a cell line (typically derived from a 129 mouse) before being bred to a relevant control strain. We genotyped our CCR9 and CXCR6 mice with the genome-wide MiniMUGAn genotyping array. We confirmed that both strains were largely (all <99.5%) of the relevant genetic background (C57BL/6NJ for CCR9 and C57BL/6J for CXCR6), with a small cluster of SNPs differing at the distal Chromosome 9 locus where the knocked-out genes are located. These SNPs were consistent with a 129 genetic background, thus confirming how these knockout strains were created.

To ensure that the presence of a 129 haplotype at the locus itself was not causal for disease differences, we identified three CC strains which contained a 129 haplotype at this locus (CC039, CC041 and CC065) and assessed the response to SARS-CoV2 in these three strains relative to CC011 and CC074 mice (**Figure S2**). While these three strains showed a range of disease responses from mild weight loss to mortality, the fact that there was not a consistent disease response in these strains suggest that the 129 haplotype at HrS26 is not informative for SARS-CoV2 disease. As such, when contrasting knockouts to relevant controls, we can be confident that the results are due to the targeted mutation itself, and not an artifact of the haplotype the mutation was generated in. Therefore, we infected *Ccr9* deficient mice (*CCR9^-/-^*) as well as the relevant age-matched C57BL/6NJ controls with SARS-CoV MA15. *CCR9^-/-^* mice showed enhanced susceptibility relative to control mice. The latter was marked by significant weight loss of >20% (HrS30), high viral load in the lungs and severe lung pathology, prolonged respiratory dysfunction (HrS26), increased lung congestion scores (HrS26), and elevated mortality (HrS30) (**Figure 5B-F**). In addition to SARS-CoV, we also studied the role of *CCR9* in clinical disease outcomes using a mouse adapted HKU3-SRBD MA (**Figure 5G-I**) (10).

We infected *CCR9^-/-^* mice and C57BL/6NJ control mice with 1×10^5^ PFU of a clade II HKU3-SRBD MA chimeric virus and monitored daily weight and other clinical measures of disease (10). *CCR9*^-/-^ mice demonstrated significantly increased weight loss compared to wild type mice beginning at 4 dpi. *CCR9*^-/-^ lungs on 4 dpi and 7 dpi also showed greater pathology than wild-type controls (p<0.005 and p<0.05, respectively). In contrast to SARS-CoV MA15 infections, HKU3-SRBD MA virus titers on 4 dpi were not significantly different between *CCR9*^-/-^ and wild type mice, with all animals clearing infectious virus by 7 dpi. These data support the hypothesis that null or low-level expression of *CCR9* likely contributes to the multitrait QTL disease phenotypes noted following clade Ia and clade II sarbecovirus infections.

Clade Ib SARS-CoV-2 MA10 infection in *CCR9^-/-^* mice also developed more severe clinical disease including greater weight loss (p<0.0001 – p<0.05) (**Figure 6A**) and virus titers (p<0.001) (**Figure 6B**) than C57BL/6NJ mice. Furthermore, infected *CCR9^-/-^* mice displayed increased PenH respiratory dysfunction (p<0.001 – p<0.01) (**Figure 6C**), lung congestion (p<0.05) (**Figure S3B**), and mortality (**Figure S3A**). Finally, *CCR9^-/-^*mice also demonstrated severe lung pathology including diffuse alveolar damage and prolonged loss of surfactant protein C (SFTPC) expression in AT2 cells, comparable to the wild-type controls (**Figure S3C-G**). Analysis of the cytokine profile in lungs by multiplex immune-assay showed increased subsets of cytokines and chemokines involved in promoting allergic airway inflammation, including IL-6, CCL3, G-CSF, CCL2, IL-13, CXCL1, and CCL11 (**Figure 6D)**. Flow cytometric analysis showed that in *CCR9*^-/-^ mice there was a significant increase in CD4^+^ T cells, CD8^+^ effector T cells, CD11^+^ dendritic cells (DCs) and eosinophils at 6 dpi, consistent with an airway inflammatory response (**Figure 6D -E**). In summary, these data support an important role of *CCR9* in protection against severe sarbecovirus disease.

**Figure 6.**
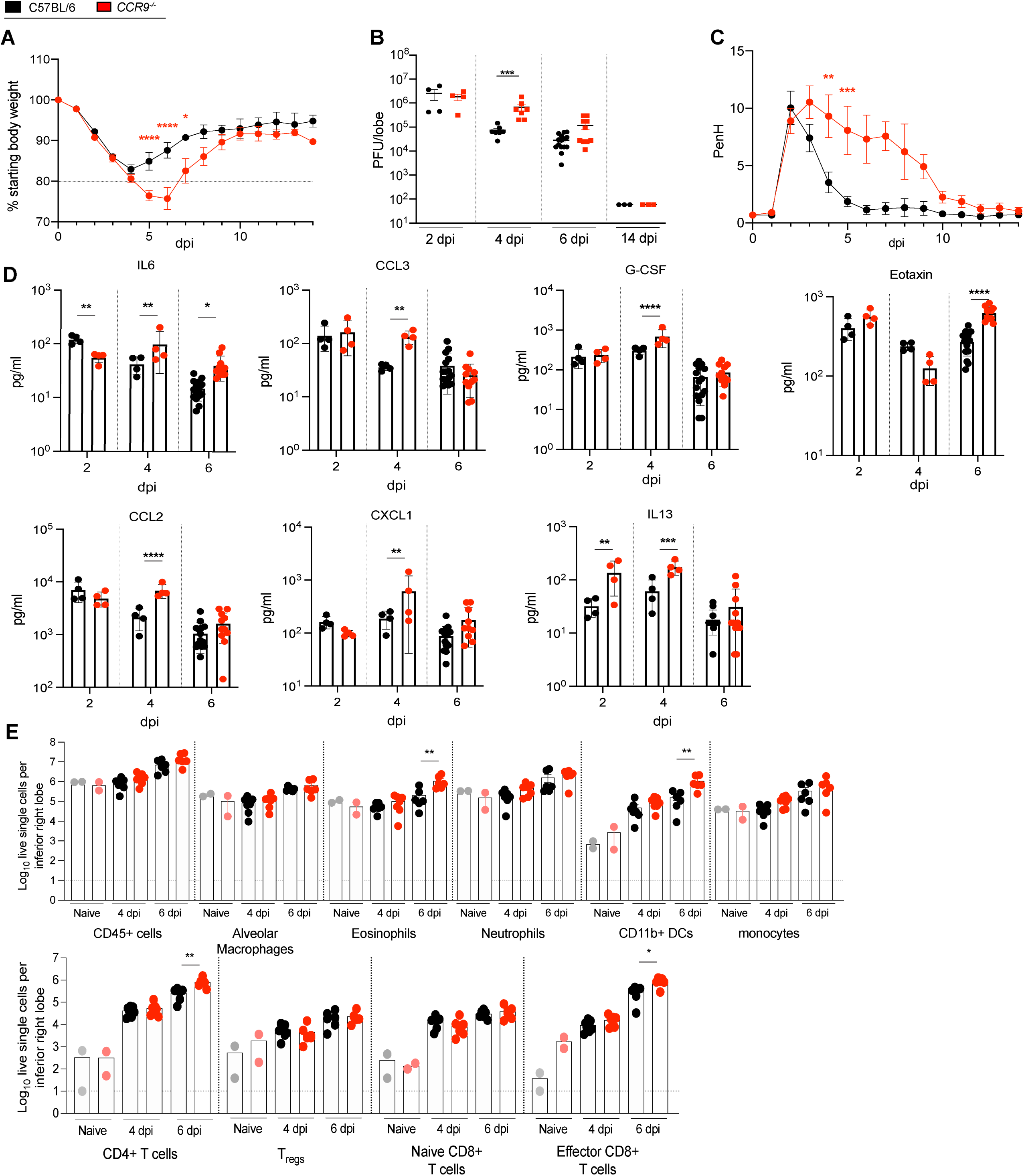
*Ccr9* Regulates Sarbecovirus Infection and Pathogenesis *in vivo*. To study the effect of *Ccr9* on the susceptibility to SARS-COV-2 infection, groups of age- matched *CCR9-/-* mice were infected with 1×10^5^ PFU SARS-CoV-2 MA10 and followed for several days for disease outcomes. A. Weight loss of *CCR9^-/-^* mice and C57BL/6NJ control mice B. Lung viral titer of *CCR9^-/-^* mice and C57BL/6NJ control mice on 2, 4, 6, and 14 dpi C. PenH of *CCR9^-/-^* mice and C57BL/6NJ control mice D. cytokine/chemokine distribution in the lung of *CCR9^-/-^* mice and C57BL/6NJ control mice on 2, 4, and 6 dpi E. Composition of lung infiltrating immune cells in the lung of *CCR9^-/-^* mice and C57BL/6NJ control mice on 4 and 6 dpi (15-18-week-old mice were infected with 1×10^5^ PFU SARS-CoV-2 MA10; C57BL/6NJ: n = 29, n = 4 for 2 dpi, n = 7 for 4 dpi, n = 15 for 6 dpi, and n = 3 for 14 dpi; *CCR9^-/-^*: n = 24 total, n = 4 for 2 dpi, n = 7 for 4 dpi, n = 10 for 6 dpi, and n = 3 for 14 dpi, respectively for weight loss, viral titer, mortality and congestion score; C57BL/6NJ: n = 4 and *CCR9^-/-^*: n = 4 lung function analysis, 57BL/6NJ: n = 4 for 2 dpi, n = 4 for 4 dpi, n = 15 for 6 dpi; *CCR9^-/-^*: n = 4 for 2 dpi, n = 4 for 4 dpi; n = 10 for 6 dpi for chemokine/cytokine analysis; n = 12 *CCR9^-/-^* and n = 13 C57BL/6NJ with n = 2 each for mock, n = 5-6 for 4 dpi, and n = 5 for 6 dpi for analysis of infiltrating cells). Data were analyzed using 2-way ANOVA with Multiple comparison (weight and PenH) and Mann-Whitney test (viral titer, congestion score, cytokine/chemokine, and infiltrating cells), *p<0.05, **p<0.005, ***p<0.0005.

### *Cxcr6* Regulates SARS-CoV MA15 and SARS-CoV-2 MA10 Pathogenesis in Mice

As the Chr9 locus (HsR26 and HsR30) also contained variants segregating in *Cxcr6* that might contribute to reduced gene expression and severe disease, we also infected *Cxcr6*- deficient (*CXCR*6^-/-^) and aged-matched controls with. SARS-CoV-MA15 and followed the mice for 7 days. We detected no significant differences in weight loss (**Figure S4B**), but significant higher viral load, congestion scores, prolonged pulmonary dysfuntion, and increased mortality (**Figures S4C-F**). The mortality rate in CXCR6-/- was significantly increased with 100% mortality by 7dpi compared to 70% in the control mice and a median survival rate of 4dpi compared to 6dpi in the control mice (**Figure S4F**). We then evaluated whether CXCR6^-/-^ also serves as a susceptibility gene for SARS-CoV-2 MA10 infection. *CXCR6-/-* mice showed more severe disease progression associated with significant weight loss (>20%) (p<0.0001 – p<0.05), prolonged airflow restriction, and increased mortality (50%) compared to wildtype C57BL/6 controls (**Figure 7A-B**) *CXCR6*^-/-^ lungs also had significantly higher viral loads and elevated congestion scores (p<0.01 for 4 dpi) (**Figure 7C-D**), compared to wild-type controls. Lung cytokine and chemokine analysis showed levels of IL-6, CCL2, G-CSF, and CXCL1 that were significantly elevated in *CXCR6^-/-^* compared to wild-type infected mice (**Figure 7E**). The composition of infiltrating cells into the lungs (**Figure 7F-G**) of *CXCR6*^-/-^ mice greater numbers of CD4^+^ T cells (**Figure 7F**), alveolar macrophages, inflammation-promoting CD11^+^ DCs and monocytes at 4 dpi (**Figure 7G**), consistent with an increased inflammatory airway response.

**Figure 7.**
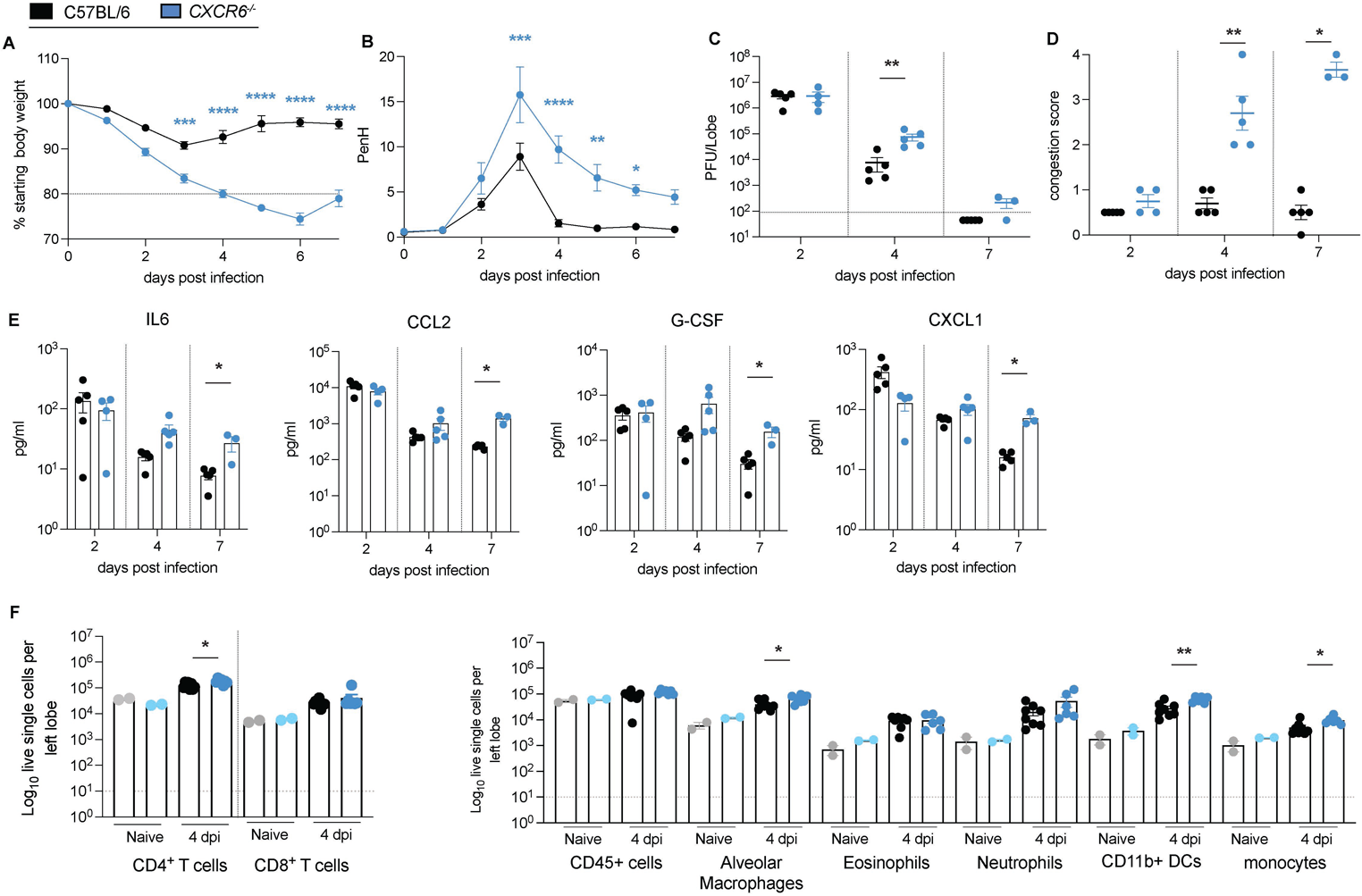
Cxcr6 Regulates SARS-CoV-2 MA10 Pathogenesis in Mice. To study the effect of *Cxcr6* on the susceptibility to SARS-COV-2 infection, groups of age- matched *CXCR6-/-* mice were infected with 1×10^5^ PFU SARS-CoV-2 MA10 and followed for several days for disease outcomes. A. Weight loss of *CXCR6^-/-^* mice and C57BL/6J control mice B. PenH of *CXCR6^-/-^* mice and C57BL/6J control mice C. Lung viral titer of *CXCR6^-/-^* mice and C57BL/6J control mice on 2, 4, and 7 dpi D. Congestion score of *CXCR6^-/-^* mice and C57BL/6J control mice on 2, 4, and 7 dpi E. Cytokine/chemokine distribution in the lung of *CXCR6^-/-^* mice and C57BL/6J control mice on 2, 4, and 7 dpi F. Composition of lung infiltrating immune cells in the lung of *CXCR6^-/-^* mice and C57BL/6J control mice on 4 dpi (15-18-week-old mice were infected with 1×10^5^ PFU SARS-CoV-2 MA10; C57BL/6NJ: n = 15, n = 5 for 2 dpi, n = 5 for 4 dpi, n = 5 for 7 dpi; *CXCR6^-/-^*: n = 13 total, n = 4 for 2 dpi, n = 5 for 4 dpi, n = 4 for 7 dpi for weight loss, viral titer, congestion score, and chemokine/cytokine analysis; C57BL/6NJ: n = 4 and *CXCR6^-/-^*: n=4 for lung function analysis, n = 8-9 *CXCR6*^-*/-*^ and n = 10 C57BL/6NJ with n = 2 each for mock, n = 6-8 for 4 dpi for analysis of infiltrating cells). Data were analyzed using 2-way ANOVA with Multiple comparison (weight and PenH) and Mann-Whitney test (viral titer, congestion score, cytokine/chemokine, and infiltrating cells); *p<0.05, **p<0.005, ***p<0.0005.

## Discussion

Emerging viruses represent an ongoing threat to human health and economic stability. As many emergence events are characterized by small outbreaks that are not amendable to GWAS, new strategies are needed to predict and understand how host genetic variation might regulate disease severity. Infectious diseases have shaped the natural genetic variation in the genomes of mammals. Although highly penetrant monogenic traits have been described that regulate disease severity in humans, most viral diseases likely are regulated by complex genetic traits (42–45). Novel viruses regularly emerge from zoonotic reservoirs to replicate and colonize one or more new mammalian hosts. Consequently, evolution may select for common susceptibility loci and genes that regulate disease severity across species, although this hypothesis remains unproven (42, 46–49). The successive emergence of genetically distinct sarbecoviruses in 2002 and 2019, which replicate in and cause severe disease in multiple mammalian hosts, provide an opportunity to determine the extent to which common or unique susceptibility loci regulate disease severity across species. During the first SARS-CoV epidemic, no consensus was reached on the human loci and genes that influence disease severity (50).

Recent GWAS in humans identified a locus 3p21.31, associated with susceptibility to respiratory failure, severe disease outcome, and hospitalizations after SARS-CoV-2 infection. This locus encompasses six genes, all of which showed strong signals in multiple GWAS studies: *SLC6A20, LZTFL1, CCR9, FYCO1, CXCR6,* and *XCR1*, with the causative signals between studies spread across these genes (14, 15, 32–34, 51). These genes are differentially regulated in mild and severe human disease, suggesting the presence of genetic variants that regulate gene expression (15, 16, 39–41). To date, none of these genes have been definitively linked to disease severity after SARS-CoV-2 infection (15, 16, 39–41). This human locus shows conserved synteny with the proximal region of our Chr9 multitrait QTL, HsR26 in mice, and shows similar genetic order and orientation of all six candidate genes. Furthermore, in addition to nonsynonymous differences in *LZTFL1*, *CCR9*, *FYCO1* and *SLC6A20A/B*, regulatory differences were seen for genes in this locus at baseline as well as in the context of infection. As such, it is likely that complex interactions between multiple candidates in this locus give rise to the observed disease differences at HrS26 and present the intriguing possibility that some of these regulatory differences underlie common disease mechanisms between mice and humans. Within the locus, we focused on two – *Ccr9* and *Cxcr6* - as both are expressed at reduced levels in our parental susceptible parental strain and in vulnerable human cases (16, 39, 40). Gene deletions represent extreme reductions in expression and loss of function, which differs from the subtle impact that allele variation in noncoding or coding regions might have. However, these studies represent an important first step towards elucidating the genetic drivers of sarbecovirus disease severity and highlight a rationale for future study of *CCR9* and *CXCR6* allele variants. *CCR9* is expressed by T cells, DCs, monocytes/macrophages and eosinophils, and is a receptor for *CCL25* (52). The *CCR9*/*CCL25* axis functions in homeostasis, inflammation and inflammation-associated diseases including early respiratory allergic inflammation, asthma, chronic inflammatory bowel diseases, and organ fibrosis (53). *CXCR6* is also expressed by T cells and serves as a homing mediator for resident memory T cells (T_RM_). Previous studies have shown that CXCR6^-/-^ mice have decreased lung resident T_RM_ cells, which is associated with increased host control of *M. tuberculosis* and influenza virus infections (54). Both *CCR9* and *CXCR6* are involved in recruiting T cells and regulating inflammatory responses during lung infection. Importantly, SARS-CoV-2 infection stimulated distinct yet overlapping infiltrating cell populations and cytokine and chemokine signatures in *CCR9^-/-^*and *CXCR6^-/-^* mice. SARS-CoV-2 infected *CCR9^-/-^*mice showed severe disease progression, with an increased influx of inflammatory macrophage/monocyte cells, T cells and eosinophils, coupled with a chemokine/cytokine profile that supports an allergic inflammatory response in the lung. SARS-CoV-2-infected *CXCR6^-/-^* mice showed an influx of inflammatory cells with expression of proinflammatory Th1-cytokines/chemokines in the lungs. As both genes are downregulated in highly susceptible CC074 mice and in humans with severe respiratory failure, it is possible that both genes, as well as other genes under the QTL, contribute to vulnerability to SARS-CoV-2 infection in mice. Detailed genetic mapping studies in human cohorts have implicated a 5’ synonymous mutation in the noncoding region of the *LZTFL1* gene, which contributes to the downregulation in expression, although it still remains uncertain if this gene directly modulates severe human or mouse disease outcomes (41).

Here, we demonstrate that mice lacking *Ccr9* or *Cxcr6 expression* developed severe disease most likely driven by excessive inflammation. The increased numbers of proinflammatory monocytes in both the *Ccr9* and *Cxcr6 deficient* mice suggest a potential mechanism for increased disease severity, as previous studies have implicated the levels of proinflammatory monocytes cells in lethal SARS-CoV infection (55). Lack of *CCR9* enhances disease severity, this gene likely has an important protective function during SARS-CoV, HKU3-SRBD, and SARS-CoV-2 infection in mice. Our data also indicates that lack of *CXCR6* gene enhances disease severity, implicating a role in protecting against lethal infection as well. In future studies, we plan to define the exact SNP(s) that regulate the differential expression levels and induction of *Ccr9* and *Cxcr6* and identify the causal mechanisms underlying the observed disease outcomes associated with the human GWAS results, as well as our *HrS26* results.

Rich and complex datasets such as described herein enable comparisons with human GWAS studies mapping QTL after SARS-CoV-2 infection. Our data supports the idea that there are shared disease susceptibility loci in both human and mice. We provide evidence from our mouse studies that the HrS26 locus is involved in disease outcomes across multiple genetically distinct sarbecoviruses. Our study highlights the power of using animal GRPs to understand the role of host genetic variation on infectious diseases, generate new models of differential disease, probe the role of individual genes in disease progression, and provide mechanistic insight into the role of specific host genes and viral strains in regulating pathogenesis across species. In appropriately selected large population screens such as the entire CC resource, highly penetrant genetic variants that impact disease outcome can be readily identified. In contrast, targeted mapping crosses between highly discordant CC RI strains can identify more complex genetic networks such as variants that are penetrant only in the context of specific genetic backgrounds, or epistatic (gene-gene) interaction networks. In addition, the multiple loci we identified, as well as unexplained heritability in this cross, suggests that CoV disease and immunity are complex polygenic traits, with the accumulation of variants across many loci driving disease susceptibility. Our studies represent a comprehensive comparison of susceptibility loci for sarbecoviruses in two different mammalian hosts, identify a large collection of susceptibility loci and candidate genes and alleles that regulate multiple aspects type-specific and cross-CoV pathogenesis, validate a potential role for the *CCR9* and/or *CXCR6* in regulating SARS-CoV-2 disease severity, and provide a resource for community-wide studies. The evidence supporting a common-susceptibility loci harbored in CC074 mice and in highly vulnerable human populations, provides new opportunities for studying sarbecovirus-host interactions, inflammation, and immunity in genetically relevant mammalian models, potentially revealing new insights into acute and chronic disease progression, and providing a potentially more relevant model for evaluating immunotherapeutic control of life threatening sarbecovirus infection outcomes.

## Acknowledgments

This study was supported by grants in aid from the National Institutes of Health, Allergy, and Infectious Diseases (AI100625 and AI149644 to R.S.B., M.T.H., M.T.F., and F.P.M.V.), a contract from the NIH (HHSN272201700036I; Task Order #38 75N93020F00001 to R.S.B), and a Burroughs Wellcome Fund Postdoctoral Enrichment Program Award and a Hanna H. Gray Fellowship from the Howard Hugues Medical Institute (DRM). We would like to thank the Systems Genetics Core Facility (UNC) for maintaining and distributing Collaborative Cross mice.

## Figure Legend

**Figure S1. Distribution of measured phenotypes and QTL maps for CC0011xCC074-F2 screen.**

A panel of 403 CC011xCC074-F2 mice were generated and infected with 1×10^4^ PFU of SARS-CoV MA15 and followed for 4 days to record clinical disease outcomes. QTL mapping was performed for several SARS-CoV-associated disease phenotypes.

A. Percentage of peripheral neutrophils of CC011xCC074-F2 mice at 4 dpi and the corresponding QTL map for the significant QTL HsR26

B. Percentage of peripheral lymphocytes of CC011xCC074-F2 mice and the corresponding QTL map for the significant QTL HsR26

C. PenH of CC011xCC074-F2 mice at 4 dpi and the corresponding QTL map for the significant QTL HsR26

D. Lung congestion score of CC011xCC074-F2 mice at 4 dpi and the corresponding QTL map for the significant QTL HsR26

E. Percentage of the overall survival of CC011xCC074-F2 mice over the time of infection and the corresponding QTL map for the significant QTL HsR26

F. Percentage of the overall survival on 4 dpi of CC011xCC074-F2 mice over the time of infection and the corresponding QTL map for the suggestive QTL HsR30

G. Percentage of starting body weight of CC011xCC074-F2 mice over the time of infection and the corresponding QTL map for the suggestive QTL HsR30

H. Percentage of starting body weight for females of CC011xCC074-F2 mice over the time of infection and the corresponding QTL map for the suggestive QTL HsR30

10-12-week-old mice were infected with 1×10^4^ PFU SARS-CoV MA15; n = 403 mice (228 male and 177 female mice); QTLs maps indicate significance of LOD at 95% (significant), 90%, and 50% (suggestive).

**Figure S2. Distribution of measured weight phenotypes CC strains with a 129 haplotype over HrS26 showed a range of disease responses.**

To validate whether a 129s1/SvImJ haplotype at HrS26 showed a disease response akin to CC074, in order to be more confident in our knockout analyses. At both day 2 (A) and day 5 (B) post infection, the three CC strains which had a 129 haplotype (CC039, CC041, CC065) showed a range of responses from ∼10% weight loss to mortality. As these responses were so variable within a haplotype, and not all strains phenocopied the susceptible CC074 response, we conclude that a 129s1 haplotype at this locus is not associated with susceptibility.(10-12-week-old mice were infected with 1×10^4^ PFU SARS-CoV-2 MA10; CC011: n=18, CC039: n=7, CC041: n=8, CC065: n=4, CC74: n=19)

**Figure S3. *CCR9^-/-^* mice show mortality and lung pathology starting at 6 dpi.**

To study the effect of *Ccr9* on the susceptibility to SARS-COV-2 infection, groups of age-matched *CCR9-/-* mice were infected with 1×10^5^ PFU SARS-CoV-2 MA10 and followed for several days for disease outcomes.

A. Survival rate of *CCR9^-/-^* mice and C57BL/6NJ control mice

B. Lung congestion score of *CCR9^-/-^*mice and C57BL/6NJ control mice on 2, 4, 6, and 14 dpi

C. Frequency (in %) of SFTPC (alveolar type 2) cells identified by RNA in situ hybridization

D. Lung pathology (Matute-Bello) of *CCR9^-/-^* mice and C57BL/6NJ control mice on 2, 4, and 6 dpi

E. Lung pathology (DAD) of *CCR9^-/-^* mice and C57BL/6NJ control mice on 2, 4, and 6 dpi

F. Representative H&E stains of lung tissue sections are shown for 4, 6, and 14 dpi for *CCR9^-/-^* and C57BL/6NJ, scale bar indicates 100μm

(15-18-week-old mice were infected with 1×10^5^ PFU SARS-CoV-2 MA10; C57BL/6NJ: n = 15, n = 5 for 2 dpi, n = 5 for 4 dpi, n = 5 for 7 dpi; *CCR9^-/-^*: n = 13 total, n = 4 for 2 dpi, n = 5 for 4 dpi, n = 4 for 7 dpi for mortality and lung congestion score; C57BL/6NJ: n = 19 with n = 4 for 2 dpi, n = 7 for 4 dpi, n = 5 for 6 dpi, n = 3 for 14 dpi; *CCR9^-/-^*: n = 20 with n = 4 for 2 dpi, n = 7 for 4 dpi, n = 6 for 6 dpi, n = 3 for 14 dpi for RNA in situ; C57BL/6NJ: n = 13 with n = 4 for 2 dpi, n = 4 for 4 dpi, n = 5 for 6 dpi; *CCR9^-/-^*: n = 10 with n = 4 for 2 dpi, n = 4 for 4 dpi, n = 2 for 6 dpi for histopathology scoring. Data were analyzed using Log-rank (mortality) and Mann-Whitney test (congestion score and pathology scores), *p<0.05, ****p<0.0001

**Figure S4. Validation of *Cxcr6* as susceptibility gene during SARS-CoV MA15 infection.**

To validate *Cxcr6* as a susceptibility gene during SARS-CoV infection, groups of age-matched *Cxcr6^-/-^* mice were infected with 1×10^5^ PFU SARS-CoV MA15 and followed for several days for disease outcomes. The Spike protein sequences of selected Sarbecoviruses were aligned and phylogenetically compared. Sequences were aligned using free end gaps with the Blosum62 cost matrix, and the tree was constructed using the neighbor-joining method with a Jukes-Cantor genetic distance model based on the multiple sequence alignment in Geneious Prime. The GenBank accession numbers for each genome sequence are shown. The tree was then output and visualized using EvolView.

A. Phylogenetic Tree of sarbecoviruses. The Spike protein sequences of selected Sarbecoviruses were aligned and phylogenetically compared. Sequences were aligned using free end gaps with the Blosum62 cost matrix, and the tree was constructed using the neighbor-joining method with a Jukes-Cantor genetic distance model based on the multiple sequence alignment in Geneious Prime. The GenBank accession numbers for each genome sequence are shown. The tree was then output and visualized using EvolView (Bold indicates viruses tested).

B. Weight loss of *Cxcr6^-/-^* mice and C57BL/6NJ control mice

C. Lung viral titer of *Cxcr6^-/-^* mice and C57BL/6NJ control mice on 4 and 6 dpi

D. PenH of *Cxcr6^-/-^* mice and C57BL/6NJ control mice

E. Congestion of *Cxcr6^-/-^* mice and C57BL/6NJ control mice on 4 and 6 dpi

F. Percentage survival of *Cxcr6^-/-^* mice and C57BL/6NJ control mice; dotted lines indicate median survival days.

(15-18-week-old mice were infected with 1×10^5^ PFU SARS-CoV-2 MA10; C57BL/6J: n = 20, n = 5 for 2 dpi, n = 5 for 4 dpi, n = 10 for 7 dpi; *CXCR6^-/-^*: n = 19 total, n = 5 for 2 dpi, n = 5 for 4 dpi, n = 9 for 7 dpi; n=4 each for lung function analysis. Data were analyzed using Log-rank (mortality) and Mann-Whitney test (congestion score and pathology scores), **p<0.005

**Figure S5. Flow cytometric analysis.** a. Flow cytometric gating strategy for lung tissue analysis.

**Table S1. Baseline gene expression of *Ccr9*, *Cxcr6*, *Xcr1*, *Lztfl1*, *Fyco1*, and *Slc6a20a/b* in selected CC strains in liver (Liv), kidney (Kid), and heart (Ht) which contain either the protective (PWK, p) or susceptible (A/J, a) haplotype at the Chr9 multitrait locus** (G. Keele, unpublished data).

## Material and Methods

### Cells and viruses

Recombinant mouse-adapted SARS-CoV MA15, HKU3-SRBD-MA (HKU3-SRBD MA), and SARS-CoV-2 MA10 virus were generated as described previously (10, 18, 35). For virus titration, the caudal lobe of the right lung was homogenized in PBS, resulting homogenate was serial-diluted and inoculated onto confluent monolayers of Vero E6 cells (ATCC CCL-81), followed by agarose overlay. Plaques were visualized with overlay of Neutral Red dye on day 2 (SARS-CoV MA15, HKU3-SRBD MA) or day 3 (SARS-CoV-2 MA10) post infection. All virological studies were conducted under BSL3 conditions, and personnel wore appropriate personal protective gear.

### Mouse studies and *in vivo* infections

Mouse studies were performed at the University of North Carolina (Animal Welfare Assurance #A3410-01) using protocols approved by the UNC Institutional Animal Care and Use Committee (IACUC). Animal studies at Washington University were carried out in accordance with the recommendations in the Guide for the Care and Use of Laboratory Animals of the National Institutes of Health. The protocols were approved by the IACUC at the Washington University School of Medicine (Assurance number A3381-01).

Mouse studies were divided into three major classes: F2 intercross mice, CC mice, and inbred wild-type or gene-edited mice. CC mice were purchased directly from the UNC Systems Genetics Core Facility at 3-5 weeks of age and acclimated for a week in the BSL3 before challenge. We contracted with the Systems Genetics Core Facility at UNC to generate the F2 mice used in this study. First, F1 mice between CC011 and CC074 were generated by cross males and females in both directions, and then the F2 mice were bred in all 4 possible F1 x F1 combinations, to ensure appropriately balanced sex Chromosome and parent-of-origin effects. F2 mice (226 males, 177 females) were weaned such that littermates were randomized to different experimental cages to reduce litter- or batch-effects on the study, and mice were transferred at 5-6 weeks of age to the laboratory for infection between 9-12 weeks of age. For studies in genetically defined knockout mice, 15-week old *CCR9^-/-^* mice (strain 027041), 15-week old *CXCR6^-/-^* mice (strain 005693) 15-week old female C57BL/6NJ mice (strain 005304), and 15-week old C57BL/6J (strain 000664) were purchased from Jackson Laboratory, and the genotype of these mutant mice were confirmed via genotyping on the MiniMUGA array (Neogen, Inc. Supplemental file X). CC mice and CC-F2 mice were inoculated with 1×10^4^ PFU (CC-F2 with SARS-CoV MA15 and CC mice with either SARS-CoV MA15 or SARS-CoV-2 MA10, respectively). *CXCR6*^-/-^, *CCR9*^-/-^, and appropriate C57BL/6 control mice were inoculated intranasally with 1×10^5^ PFU of either SARS-CoV MA15, SARS-CoV-2 MA10, or HKU3-SRBD MA in 50 μl of PBS. Body weight, mortality, and pulmonary function by whole body plethysmography (56) were monitored daily as indicated. At the designated timepoints, mice were euthanized and gross pathology (congestion score) of the lung was assessed and scored on a scale from 0 (no lung congestion) to 4 (severe congestion affecting all lung lobes). Then, lung tissue was harvested for titer and histopathology analysis, and blood samples were harvested for analysis of immune cells. Samples were stored at -80°C until homogenized and titered by plaque assay as described above. Peripheral blood was diluted 1:5 in PBS/EDTA and analyzed with the VetScan HM5 as previously described (57). Histopathology samples were fixed in 10% phosphate buffered formalin for 7 days before paraffin embedding, sectioning stained with hematoxylin and eosin and scored for DAD and acute lung injury as previously described by our group (58).

### Flow cytometry analysis of immune cell infiltrates

For analysis of lung tissues, mice were perfused with sterile PBS, and the right inferior lung lobes were digested at 37°C with 630 µg/mL collagenase D (Roche) and 75 U/mL of DNase I (Sigma–Aldrich) for 2 h. Single cell suspensions were preincubated with Fc Block antibody (BD PharMingen) in PBS + 2% heat-inactivated FBS for 10 min at room temperature before staining. Cells were incubated with antibodies against the following markers: efluor506 Viability Dye (Thermo Fisher, 65-0866-14), BUV395 anti-CD45 (Clone 30-F11, BD Biosciences), BV711 anti-CD11b (Clone M1/70, Biolegend), APC-Cy7 anti-CD11c (Clone HL3, BD Biosciences), BV650 anti-Ly6G (Clone 1A8, Biolegend), Pacific Blue anti-Ly6C (Clone HK1.4, Biolegend) FITC anti-CD24 (Clone M1/69, Biolegend), PE anti-Siglec F (Clone E50-2440, Biolegend), PE-Cy7 anti-CD64 (Clone X54-5/7.1, Biolegend), AF700 anti-MHCII (Clone M5/114.15.2, Biolegend), BV421 anti-CD3 (Clone 17A2, Biolegend), BV785 anti-CD4 (Clone GK1.5, Biolegend), APC anti-CD8a (Clone 53-6.7, Biolegend) BV421 anti-B220 (Clone RA3-6B2, Biolegend) APC-Cy7 anti-CD44 (Clone IM7, Biolegend) BV605 anti-CD62L (Clone MEL-14, Biolegend). All antibodies were used at a dilution of 1:200. Cells were stained for 20 min at 4°C, washed, fixed and permeabilized for intracellular staining with Foxp3/Transcription Factor Staining Buffer Set (eBioscience) according to manufacturer’s instructions. Cells were incubated overnight at 4°C with BV421 anti-Foxp3 (Clone MF-14, Biolegend) washed, re-fixed with 4% PFA (EMS) for 20 min and resuspended in permeabilization buffer. Absolute cell counts were determined using Trucount beads (BD). Data were acquired on a flow cytometer (BD-X20; BD Biosciences) and analyzed using FlowJo software (Tree Star) (**Figure S5**).

### Cytokine and chemokine protein analysis

The small center lung lobe of each mouse was homogenized in 1 ml of PBS and briefly centrifuged to remove debris. Fifty microliters of homogenate were used to measure cytokine and chemokine protein abundance using a Bio-Plex Pro mouse cytokine 23-plex assay (Bio-Rad) according to the manufacturer’s instructions.

### Lung pathology scoring and RNA in situ hybridization/quantification

Two separate lung pathology scoring scales, Matute-Bello and Diffuse Alveolar Damage (DAD), were used to quantify acute lung injury (58). For Matute-Bello scoring samples were blinded and three random fields of lung tissue were chosen and scored for the following: (A) neutrophils in alveolar space (none = 0, 1–5 cells = 1, > 5 cells = 2), (B) neutrophils in interstitial space (none = 0, 1–5 cells = 1, > 5 cells = 2), (C) hyaline membranes (none = 0, one membrane = 1, > 1 membrane = 2), (D) Proteinaceous debris in air spaces (none = 0, one instance = 1, > 1 instance = 2), (E) alveolar septal thickening (< 2Å∼ mock thickness = 0, 2–4Å∼ mock thickness = 1, > 4Å∼ mock thickness = 2). Scores from A–E were put into the following formula score = [(20x A) + (14 x B) + (7 x C) + (7 x D) + (2 x E)]/100 to obtain a lung injury score per field and then averaged for the final score for that sample.

In a similar manner, DAD scoring was obtained by evaluating three random fields of lung tissue that were scored for the in a blinded manner for: 1= absence of cellular sloughing and necrosis, 2= uncommon solitary cell sloughing and necrosis (1–2 foci/field), 3=multifocal (3+foci) cellular sloughing and necrosis with uncommon septal wall hyalinization, or 4=multifocal ( >75% of field) cellular sloughing and necrosis with common and/or prominent hyaline membranes. To obtain the final DAD score per mouse, the scores for the three fields per mouse were averaged.

RNA-ISH was performed on paraffin-embedded 5 μm tissue sections using the RNAscope Multiplex Fluorescent Assay v2 according to the manufacturer’s instructions (Advanced Cell Diagnostics). Briefly, tissue sections were deparaffinized with xylene and 100% ethanol twice for 5min and 1 min, respectively, incubated with hydrogen peroxide for 10 min and in boiling water for 15 min, and then incubated with Protease Plus (Advanced Cell Diagnostics) for 15 min at 40°C. Slides were hybridized with custom probe (RNAscope® Probe-Mm-Sftpc-C2, cat no. 314101-C2) at 40°C for 2 h, and signals were amplified according to the manufacturer’s instructions. Stained sections were scanned and digitized by using an Olympus VS200 fluorescent microscope with a 40X 1.35 NA objective. Images were imported into Visiopharm Software® (version 2020.09.0.8195) for quantification. Lung tissue and Sftpc signal were quantified using a customized analysis protocol package to 1) detect lung tissue using a decision forest classifier, 2) detect the probe signal based on the intensity of the signal in the channel corresponding to the relevant probe (Sftpc). All slides were analyzed under the same conditions. Results were expressed as the area of the probe relative to total lung tissue area.

### Mouse DNA Genotyping

CC011, CC074, their F1 progeny, the F2 cross, CXCR6^-/-^, CCR9^-/-^ mice, and appropriate controls were genotyped on the MiniMUGA genotyping array (28, 37). Genomic DNA was isolated from tail-clips of animals using the Qiagen (Hilden, Germany) DNeasy Blood & Tissue kit. 1.5 μg was sent to Neogen (Lincoln, Nebraska) for processing. We filtered the genotypes upon return for informativeness within this cross. To be considered as valid, the marker had to have one homozygous allele in all CC011 mice genotyped, the alternate homozygous allele in all CC074 mice genotyped, and the appropriate call in all F1 animals (H calls on the autosomes, an H call in females on the X Chromosomes, and the relevant homozygous call in male F1s). This filtering reduced the ∼10,800 SNPs on the MiniMUGA array to 2821 informative markers.

### QTL mapping and statistical analyses

We used R/QTL for genetic mapping (59, 60). Briefly, after appropriate data transformations, we used the scanone function to determine the strength of the regression of phenotypes on genotypes at each of the informative markers in the cross. Significance thresholds were determined by running 1000 permutations, scrambling the relationship between phenotypes and genotypes, providing an appropriate threshold of significance that is robust to the phenotype distribution and allele frequencies. For phenotypes where we identified multiple QTL, we ensured that long-range linkage disequilibrium was not driving these observations by fitting multi-factor ANOVAs with single QTL and with sets of loci. Only loci for which there was a statistically significant improvement in fit for the full model were kept.

### Data availability

All data supporting the finding of this study are available within the manuscript and are available from the corresponding authors upon request.

### Statistical analysis

The number of independent experiments and technical replicates are indicated in the relevant figure legends. Two-way ANOVA with multiple comparison were used for weight loss and lung function comparisons, log-rank was used for survival studies, and Mann-Whitney test was used for viral titer, congestion score, okine/cytokine, infilatrating cells, histophatholgy score, gene expression comparisons.

### Code use

We used R/QTL for genetic mapping and the scanone function to determine the strength of the regression of phenotypes on genotypes at each of the informative markers in the cross.

## References

1. D. Ge et al., Genetic variation in IL28B predicts hepatitis C treatment-induced viral clearance. Nature 461, 399–401 (2009).

2. P. J. McLaren et al., Polymorphisms of large effect explain the majority of the host genetic contribution to variation of HIV-1 virus load. Proc Natl Acad Sci U S A 112, 14658–14663 (2015).

3. B. Chen et al., Overview of lethal human coronaviruses. Signal Transduct Target Ther 5, 89 (2020).

4. Q. Wang, A. N. Vlasova, S. P. Kenney, L. J. Saif, Emerging and re-emerging coronaviruses in pigs. Curr Opin Virol 34, 39–49 (2019).

5. Y. Z. Zhang, E. C. Holmes, A Genomic Perspective on the Origin and Emergence of SARS-CoV-2. Cell 181, 223–227 (2020).

6. P. Zhou et al., A pneumonia outbreak associated with a new coronavirus of probable bat origin. Nature 579, 270–273 (2020).

7. V. D. Menachery et al., SARS-like WIV1-CoV poised for human emergence. Proc Natl Acad Sci U S A 113, 3048–3053 (2016).

8. S. J. Anthony et al., Further Evidence for Bats as the Evolutionary Source of Middle East Respiratory Syndrome Coronavirus. mBio 8 (2017).

9. V. D. Menachery et al., A SARS-like cluster of circulating bat coronaviruses shows potential for human emergence. Nat Med 21, 1508–1513 (2015).

10. M. M. Becker et al., Synthetic recombinant bat SARS-like coronavirus is infectious in cultured cells and in mice. Proc Natl Acad Sci U S A 105, 19944–19949 (2008).

11. M. Enserink, K. Kupferschmidt, With COVID-19, modeling takes on life and death importance. Science 367, 1414–1415 (2020).

12. T. Ahmad, Haroon, M. Baig, J. Hui, Coronavirus Disease 2019 (COVID-19) Pandemic and Economic Impact. Pak J Med Sci 36, S73–S78 (2020).

13. C. D. Eckstrand et al., An outbreak of SARS-CoV-2 with high mortality in mink (Neovison vison) on multiple Utah farms. PLoS Pathog 17, e1009952 (2021).

14. D. Ellinghaus et al., Genomewide Association Study of Severe Covid-19 with Respiratory Failure. N Engl J Med 10.1056/NEJMoa2020283 (2020).

15. E. Pairo-Castineira et al., Genetic mechanisms of critical illness in Covid-19. Nature 10.1038/s41586-020-03065-y (2020).

16. Y. Dai et al., Association of CXCR6 with COVID-19 severity: delineating the host genetic factors in transcriptomic regulation. Hum Genet 140, 1313–1328 (2021).

17. Q. Zhang et al., Inborn errors of type I IFN immunity in patients with life-threatening COVID-19. Science 370 (2020).

18. S. R. Leist et al., A Mouse-adapted SARS-CoV-2 induces Acute Lung Injury (ALI) and mortality in Standard Laboratory Mice. Cell 10.1016/j.cell.2020.09.050 (2020).

19. D. G. Ashbrook et al., A platform for experimental precision medicine: The extended BXD mouse family. Cell Syst 12, 235–247 e239 (2021).

20. J. B. Graham et al., Baseline T cell immune phenotypes predict virologic and disease control upon SARS-CoV infection in Collaborative Cross mice. PLoS Pathog 17, e1009287 (2021).

21. A. Srivastava et al., Genomes of the Mouse Collaborative Cross. Genetics 206, 537–556 (2017).

22. J. R. Shorter et al., Whole Genome Sequencing and Progress Toward Full Inbreeding of the Mouse Collaborative Cross Population. G3 (Bethesda) 9, 1303–1311 (2019).

23. S. R. Leist, R. S. Baric, Giving the Genes a Shuffle: Using Natural Variation to Understand Host Genetic Contributions to Viral Infections. Trends Genet 34, 777–789 (2018).

24. A. Schafer, R. S. Baric, M. T. Ferris, Systems approaches to Coronavirus pathogenesis. Curr Opin Virol 6, 61–69 (2014).

25. M. T. Ferris et al., Modeling host genetic regulation of influenza pathogenesis in the collaborative cross. PLoS Pathog 9, e1003196 (2013).

26. J. B. Graham et al., Immune Predictors of Mortality After Ribonucleic Acid Virus Infection. J Infect Dis 221, 882–889 (2020).

27. L. E. Gralinski et al., Genome Wide Identification of SARS-CoV Susceptibility Loci Using the Collaborative Cross. PLoS Genet 11, e1005504 (2015).

28. L. E. Gralinski et al., Allelic Variation in the Toll-Like Receptor Adaptor Protein Ticam2 Contributes to SARS-Coronavirus Pathogenesis in Mice. G3 (Bethesda) 7, 1653–1663 (2017).

29. K. E. Noll et al., Complex Genetic Architecture Underlies Regulation of Influenza-A-Virus-Specific Antibody Responses in the Collaborative Cross. Cell Rep 31, 107587 (2020).

30. A. L. Rasmussen et al., Host genetic diversity enables Ebola hemorrhagic fever pathogenesis and resistance. Science 346, 987–991 (2014).

31. P. L. Maurizio et al., Bayesian Diallel Analysis Reveals Mx1-Dependent and Mx1-Independent Effects on Response to Influenza A Virus in Mice. G3 (Bethesda) 8, 427–445 (2018).

32. J. F. Shelton et al., Trans-ancestry analysis reveals genetic and nongenetic associations with COVID-19 susceptibility and severity. Nat Genet 10.1038/s41588-021-00854-7 (2021).

33. C.-H. G. Initiative, Mapping the human genetic architecture of COVID-19. Nature 10.1038/s41586-021-03767-x (2021).

34. G. H. L. Roberts, et al., AncestryDNA COVID-19 Host Genetic Study Identifies Three Novel Loci. medRxiv 10.1101/2020.10.06.20205864, 2020.2010.2006.20205864 (2020).

35. A. Roberts et al., A mouse-adapted SARS-coronavirus causes disease and mortality in BALB/c mice. PLoS Pathog 3, e5 (2007).

36. K. W. Broman et al., R/qtl2: Software for Mapping Quantitative Trait Loci with High-Dimensional Data and Multiparent Populations. Genetics 211, 495–502 (2019).

37. J. S. Sigmon et al., Content and Performance of the MiniMUGA Genotyping Array: A New Tool To Improve Rigor and Reproducibility in Mouse Research. Genetics 216, 905–930 (2020).

38. W. S. Institute (https://www.sanger.ac.uk/sanger/Mouse_SnpViewer/rel-1505.

39. S. Kasela et al., Integrative approach identifies SLC6A20 and CXCR6 as putative causal genes for the COVID-19 GWAS signal in the 3p21.31 locus. Genome Biol 22, 242 (2021).

40. Y. Yao et al., Genome and epigenome editing identify CCR9 and SLC6A20 as target genes at the 3p21.31 locus associated with severe COVID-19. Signal Transduct Target Ther 6, 85 (2021).

41. D. J. Downes et al., Identification of LZTFL1 as a candidate effector gene at a COVID-19 risk locus. Nat Genet 53, 1606–1615 (2021).

42. L. Lindesmith et al., Human susceptibility and resistance to Norwalk virus infection. Nat Med 9, 548–553 (2003).

43. T. Dragic et al., HIV-1 entry into CD4+ cells is mediated by the chemokine receptor CC-CKR-5. Nature 381, 667–673 (1996).

44. Y. Huang et al., The role of a mutant CCR5 allele in HIV-1 transmission and disease progression. Nat Med 2, 1240–1243 (1996).

45. M. A. Jimenez-Sousa et al., IL28RA polymorphism is associated with early hepatitis C virus (HCV) treatment failure in human immunodeficiency virus-/HCV-coinfected patients. J Viral Hepat 20, 358–366 (2013).

46. R. Green et al., Oas1b-dependent Immune Transcriptional Profiles of West Nile Virus Infection in the Collaborative Cross. G3 (Bethesda) 7, 1665–1682 (2017).

47. P. Horby, N. Y. Nguyen, S. J. Dunstan, J. K. Baillie, The role of host genetics in susceptibility to influenza: a systematic review. PLoS One 7, e33180 (2012).

48. A. Kambhampati, D. C. Payne, V. Costantini, B. A. Lopman, Host Genetic Susceptibility to Enteric Viruses: A Systematic Review and Metaanalysis. Clin Infect Dis 62, 11–18 (2016).

49. J. K. Lim et al., Genetic variation in OAS1 is a risk factor for initial infection with West Nile virus in man. PLoS Pathog 5, e1000321 (2009).

50. W. K. Ip et al., Mannose-binding lectin in severe acute respiratory syndrome coronavirus infection. J Infect Dis 191, 1697–1704 (2005).

51. T. P. Velavan et al., Host genetic factors determining COVID-19 susceptibility and severity. EBioMedicine 72, 103629 (2021).

52. S. Uehara, A. Grinberg, J. M. Farber, P. E. Love, A role for CCR9 in T lymphocyte development and migration. J Immunol 168, 2811–2819 (2002).

53. X. Wu et al., The Roles of CCR9/CCL25 in Inflammation and Inflammation-Associated Diseases. Front Cell Dev Biol 9, 686548 (2021).

54. A. S. Ashhurst et al., CXCR6-Deficiency Improves the Control of Pulmonary Mycobacterium tuberculosis and Influenza Infection Independent of T-Lymphocyte Recruitment to the Lungs. Front Immunol 10, 339 (2019).

55. R. Channappanavar et al., Dysregulated Type I Interferon and Inflammatory Monocyte-Macrophage Responses Cause Lethal Pneumonia in SARS-CoV-Infected Mice. Cell Host Microbe 19, 181–193 (2016).

56. V. D. Menachery, L. E. Gralinski, R. S. Baric, M. T. Ferris, New Metrics for Evaluating Viral Respiratory Pathogenesis. PLoS One 10, e0131451 (2015).

57. S. R. Leist, K. L. Jensen, R. S. Baric, T. P. Sheahan, Increasing the translation of mouse models of MERS coronavirus pathogenesis through kinetic hematological analysis. PLoS One 14, e0220126 (2019).

58. T. P. Sheahan et al., Comparative therapeutic efficacy of remdesivir and combination lopinavir, ritonavir, and interferon beta against MERS-CoV. Nat Commun 11, 222 (2020).

59. D. M. Gatti et al., Quantitative trait locus mapping methods for diversity outbred mice. G3 (Bethesda) 4, 1623–1633 (2014).

60. K. W. Broman, H. Wu, S. Sen, G. A. Churchill, R/qtl: QTL mapping in experimental crosses. Bioinformatics 19, 889–890 (2003).

